# Beyond the average: An updated framework for understanding the relationship between cell growth, DNA replication, and division in a bacterial system

**DOI:** 10.1101/2022.03.15.484524

**Authors:** Sara Sanders, Kunaal Joshi, Petra Anne Levin, Srividya Iyer-Biswas

## Abstract

Our current understanding of the bacterial cell cycle is framed largely by population-based experiments that focus on the behavior of idealized average cells. Most famously, the contributions of Cooper and Helmstetter help to contextualize the phenomenon of overlapping replication cycles observed in rapidly growing bacteria. Despite the undeniable value of these approaches, their necessary reliance on the behavior of idealized average cells washes out the stochasticity inherent in single cell growth and physiology limiting their mechanistic value. To bridge this gap, we propose an updated and agnostic framework, informed by extant single-cell data, that quantitatively accounts for stochastic variations in single-cell dynamics and the impact of medium composition on cell growth and cell cycle progression. In this framework, stochastic timers sensitive to medium composition impact the relationship between cell cycle events, accounting for observed differences in the relationship between cell cycle events in slow and fast growing cells. We conclude with a roadmap for potential application of this framework to longstanding open questions in the bacterial cell cycle field.

## Introduction

Proliferation of organisms across the tree of life requires effective coordination of cell growth, DNA replication, and division. Coordination is challenging in bacteria, for which population mass doubling times (“MDTs”) can vary as much as five-fold with nutrient availability. In many bacteria, including the model organism *Escherichia coli,* the time required to complete a round of DNA replication can be longer than the MDT, particularly under nutrient rich conditions, resulting in multiple ongoing cycles of DNA replication on the same chromosomal template.^1,2^

Until recently, bacteriologists relied on population-level strategies to understand the relationship between cell growth and cell cycle progression. These approaches wash out stochastic, cell-to-cell variations, resulting in models that do not always hold up in single cells.^3–9^ In particular, the MDT or “growth rate” is frequently misinterpreted as a determinant variable that directly impacts the rate of cell-cycle progression, to the contrary of the original phenomenological description by Helmstetter and Cooper.^1,10^ Relatively recent advances in microfluidics and single-cell analysis are beginning to reveal the principles governing bacterial cell-cycle progression at the single-cell level. Here, we review the prevailing population-based models of bacterial growth and cell cycle progression, highlighting the core reasoning underlying each. Next, we leverage extant data to propose a framework from which to understand the bacterial cell cycle accounting for physiology and stochasticity inherent in single cells. Finally, we end with a discussion of open questions and avenues for future research.

## The nutrient growth law

Early work on the *E. coli* cell cycle focused on the relationships between growth rate and four parameters: cell size, RNA content, DNA content, and nutrient composition. In their classic 1958 study, Schaechter, Maaløe, and Kjeldgaard (SMK) observed that *Salmonella* Typhimurium cell mass increases exponentially with nutrient-imposed mass doubling time (MDT).^11^ Protein, RNA, and DNA content similarly increase, indicating that their overall concentration (mass/mass) remains constant.

Although their data is noisy (see Figure 1 of SMK for example related to size), the best-fit line drawn through 22 different media conditions suggests a simplified model in which each of the four measured parameters (size, protein, DNA, and RNA content) depend on growth rate. “At a given temperature, size and composition are found to depend in a simple manner on the growth rate afforded by the medium. This *implies* that media that give identical growth rates produce identical physiological states, *regardless of the actual constituents of the media*”^11^ (emphasis added). Based on analysis of cells cultured at different temperatures, SMK further clarify that culture medium composition dictates growth rate, and it ultimately dictates the chemical composition of the cell. Despite this important caveat, growth rate—not medium composition—quickly became perceived as the primary driver of cell cycle progression.^12^ In part, because of the simplicity with which it lends itself to mathematical modeling. This positive relationship between growth rate, cellular composition, and cell cycle progression is colloquially referred to as the “growth law” or “nutrient growth law”.^11,13,14^

**Fig. 1:**
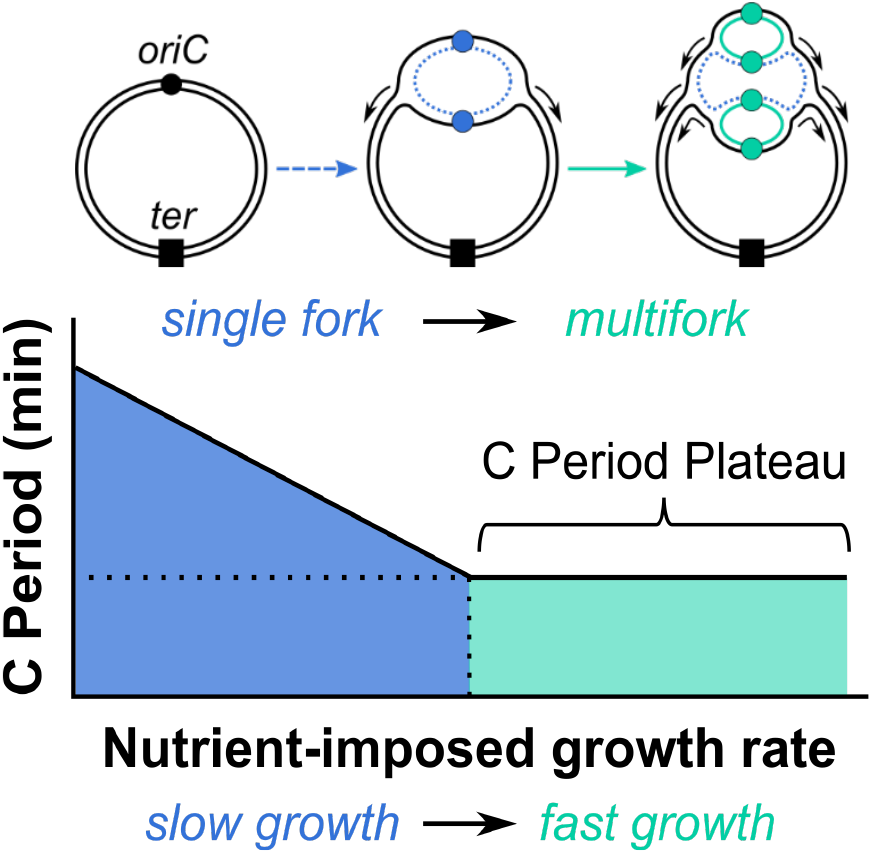
Replication of the *E. coli* chromosome. *Top:* Bacterial chromosome depicting the origin of replication *(oriC,* •) and terminus *(ter,* ■). After birth, replication initiation yields a single replication bubble, a replication state termed “single fork replication” 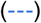. When a second initiation event occurs before termination of the prior round, “multifork replication” occurs 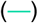. Active origins and newly synthesized DNA are indicated with colors corresponding to replication state. *Bottom:* In the Cooper-Helmstetter model, as nutrient-imposed growth rate increases, the C period length decreases until it reaches a plateau during fast growth.

## The Cooper-Helmstetter model of cell cycle progression

Once SMK identified a positive connection between nutrient-imposed growth rate and composition, the next challenge was to determine how this connection was achieved. Focusing on DNA replication and leveraging their ability to synchronize cells with their “baby machine,” Cooper and Helmstetter analyzed DNA synthesis in *E. coli* in real-time across 13 different media compositions.^15^

Incorporating SMK’s findings^11^ and work from Cairns^16^ identifying the circular nature of the bacterial chromosome, Cooper and Helmstetter developed a quantitative phenomenological model of the bacterial cell cycle (the “CH model”).^1^ The CH model posits two distinct replication regimes: single fork and multifork. Single fork (really single round) replication with up to 2 forks proceeding at a time prevails during slow growth, and the time required to complete a round of DNA replication (C-period) varies with nutrient-imposed population growth rate as does the period between the initiation of new rounds of DNA replication and the initiation of new rounds of cell division.^1,15^ During the multifork regime, division and initiation continue to co-vary with population mass doubling time, however C period becomes fixed.^1^ (**Fig. 1**) Because of this imbalance, new rounds of replication are started prior to completion of the previous one providing an explanation for the multiple origins of replication observed by Cooper and Helmstetter and others,^1,15,17–22^ including later studies that assess cell size.^8,12,23–26^

## Population and single cell data tell different stories

As Cooper and Helmstetter themselves note, their model applies specifically to idealized average cells. They intended to explain the phenomena of multifork replication as overlapping replication cycles, not to provide a mechanistic framework from which to understand relationships between cell cycle events.^10^ In individual cells, stochasticity adds another layer of complexity to the already inherently complex process of overlapping cell cycles. Indeed, single-cell data reveal high levels of stochasticity regarding both growth rate and the temporal progression of cell cycle events. (**Fig. 2**) Even when the mass doubling time of a population is held constant, the instantaneous growth rate of single-cells within that population varies as much as three-fold.^8,27,28^ Additionally, while single-cell data support a diminishing relationship between growth rate and replisome speed in faster-growing subpopulations.^8,13,27,28^ *E. coli* elongation rates vary as much as fourfold (250 to 1000 nucleotides/second, nt/s).^29–31^ Finally, the “nutrient growth law” proposed by SMK is inextricably linked to the CH model, despite accumulating evidence that growth rate varies independent of cell size and cell cycle progression, although all three are sensitive to nutrient availability and quality.^11,32–34^ DNA replication is inherently sensitive to medium composition as it directly impacts the availability of nucleotide precursors, and a coterie of mutations are known to impact size and/or DNA replication-independent of nutrient-imposed growth rate.^32,34–37^

**Fig. 2:**
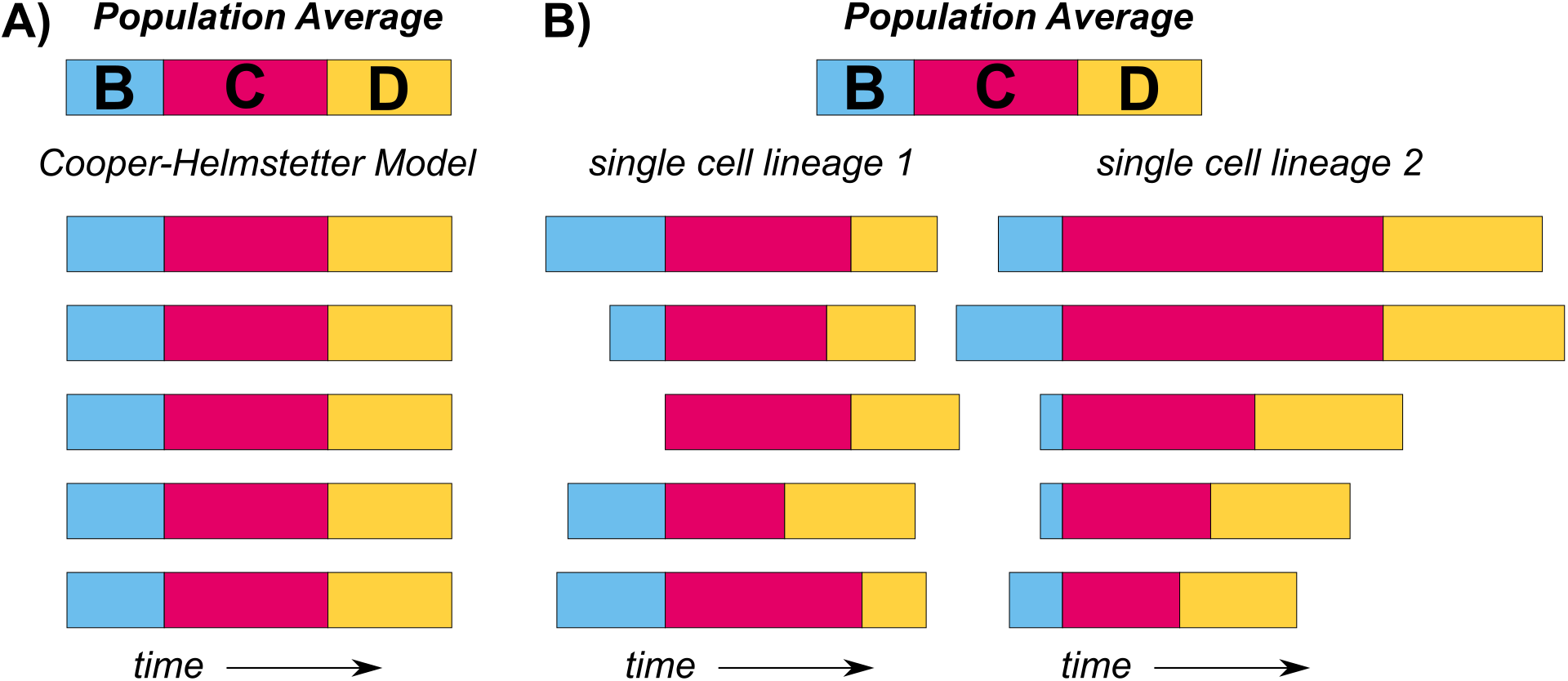
Deterministic paradigm vs. stochastic nature of cell cycle timescales. (A) The CH model assumes that all cells within a given condition follow the population average B, C, and D periods. (B) Relative B, C, and D periods are shown over multiple consecutive replication cycles for two cell lineages grown on MOPS glucose (based on data from Si et al.). Significant differences between replication cycles necessitate a new theory accounting for stochasticity.

## An agnostic framework leveraging stochastic timers illuminates the relationship between nutrient composition and cell cycle progression in individual cells

The disconnects outlined above—the inability to account for stochasticity and the misassumption that growth rate is a primary driver of cellular physiology rather than medium composition— highlight the deficiencies of the CH model as a universal tool for understanding the mechanisms underlying bacterial cell cycle control. To address this gap, we leveraged published single-cell datasets for slow and intermediate growth regimes^8^ to develop an agnostic framework for medium-dependent stochastic bacterial cell cycle progression.

Addressing all problems listed above at once, our framework centers on the idea that at each initiation event, three new stochastic “timers”—corresponding respectively to single-cell inter-initiation time, *τ_*i*_*; fork completion time (C period duration of individual cells), *τ_c_*; and the time between initiation and the corresponding division event (C+D period for individual cells), *τ_d_*—all begin to tick (**Fig. 3**). Thus, the relative order of completion of these three timers determines the fork number at the start of the next replication cycle. We elected to begin with initiation rather than other cell cycle events as this step is traditionally viewed as the beginning of the bacterial cell cycle. In *E. coli,* replication initiation is tied to cell growth via accumulation of the initiator protein DnaA to threshold levels.^8,38–43^

**Fig. 3:**
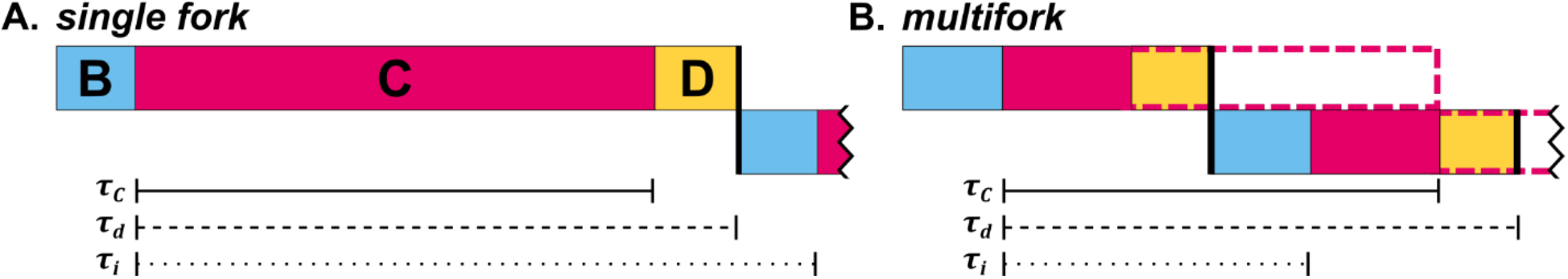
Stochastic timers. Visual representations of the timers *τ_i_, τ_c_* and *τ_d_* in a cell undergoing single fork (A) and multifork (B) replication. These stochastic timers represent single-cell parameters. During multifork replication, C periods extend beyond a single division cycle. This overlap is indicated by extended dotted lines. Color schemes match Fig. 2.

We observed that the distributions of the three single-cell variables (*τ_i_*, *τ_c_* and *τ_d_*) generally vary with the stochastic single-cell exponential growth rate, *k,* consistent with previous single cell data.^8,27,28^ Thus, we extract the stochastic distributions of these timers along with the subsequent Thus, we extract the stochastic distributions of these timers along with the subsequent replication cycle’s growth rate as experimentally measured functions of *k* (see supplementary section for details), different for each independent growth condition.

To account for media dependent variations in timer distributions, we generated “calibration curves” for each media condition and used these curved as input in our framework to simulate the next *τ_i_*, *τ_c_*, *τ_d_*, and *k* for each consecutive replication cycle. (**Fig. S1**) This framework is agnostic to the specific mechanisms governing the dependence on *k,* and thus robust to nutrient-dependent or strain-dependent differences in cell growth and cell cycle progression.

Under slow growth conditions (M9 acetate), cells typically initiate and complete a single round of replication (single fork) per cell cycle (2-fork cycle in **Fig. 4**, CH model slow growth regime).^1,15,44^ Mathematically, the hierarchy between the three timers can be described as *τ_i_* > *τ_d_* > *τ_c_*. However, in some cases, after termination we observe a new round of replication initiation prior to division (*τ_d_* > *τ_i_* > *τ_c_*), leading to a state with a total of four active replication forks divided across two chromosomes *(ori:ter* = 4:2, **Fig. 4**). Thus, the number of replication forks per cell immediately after initiation depends only on the order of division relative to initiation. Altogether, in slow growth conditions, our framework can be simplified to require input of only two timers as functions of *k*: (i) **inter-initiation time**, *τ_i_*; and (ii) **time to division**, *τ_d_*. Using this streamlined version and the “calibration curves” obtained from the same single-cell dataset (**Fig. S1**), we simulate the population distribution of the number of replication forks per cell. The results of our fitting parameter-free simulations match experimental data extremely well, thus validating the soundness of the conceptual framework (**Fig. 5A**).

**Fig. 4:**
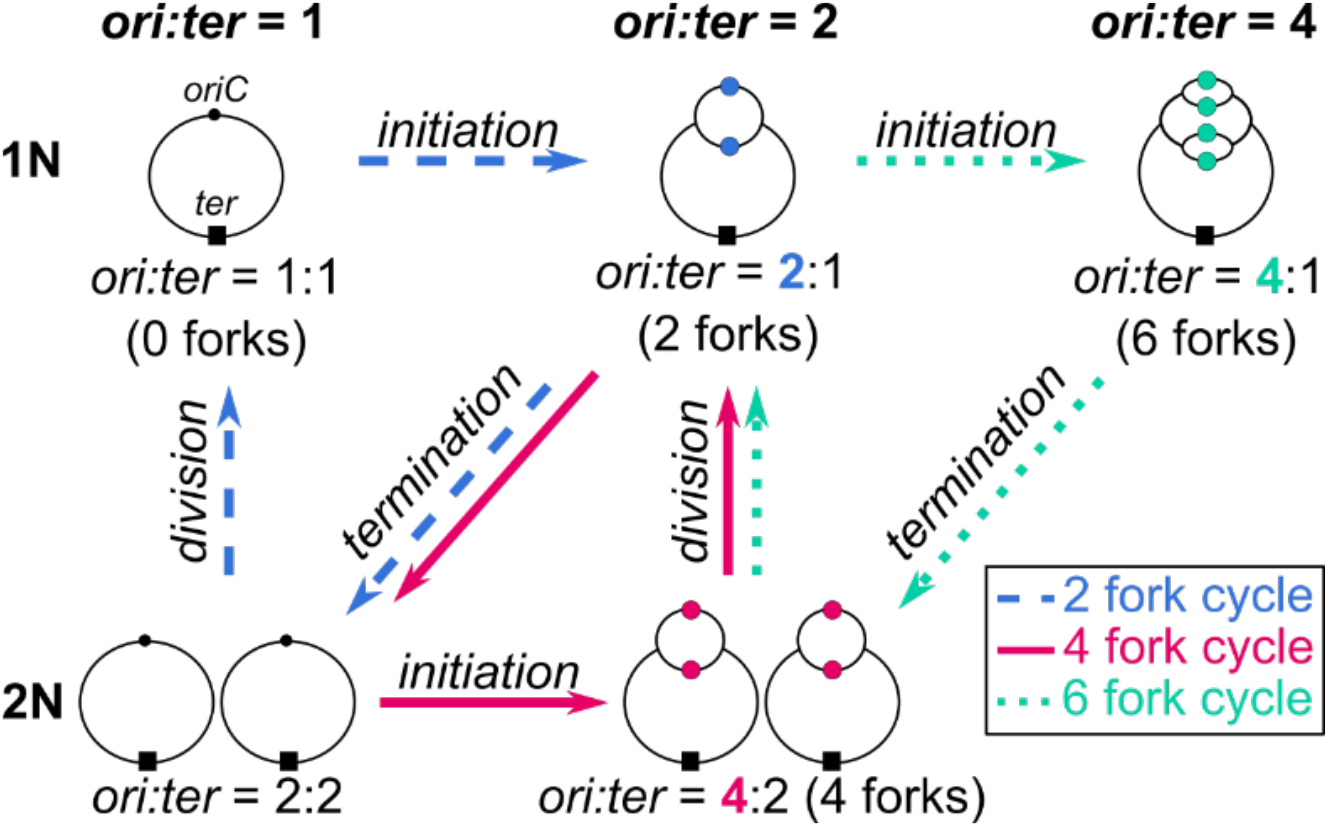
Flow chart depicting the possible replication cycles depending on the order of initiation, termination, and division events. *N* represents the number of chromosome copies present in the cell and is equal to the number of termini *(ter,* ■). 2-fork cycle (blue) forms 2 forks upon initiation at *oriC,* •, then replication is completed leaving 0 forks. 4-fork cycle (magenta) progresses from 4 forks at initiation to 2 forks after division to termination. 6-fork cycle (teal) progresses from 6 forks at initiation to 4 forks after termination of previous replication to 2 forks after division to 6 forks after new round of initiation. Further configurations with 8 forks or more are relevant only at growth rates faster than available in this dataset.

**Fig. 5:**
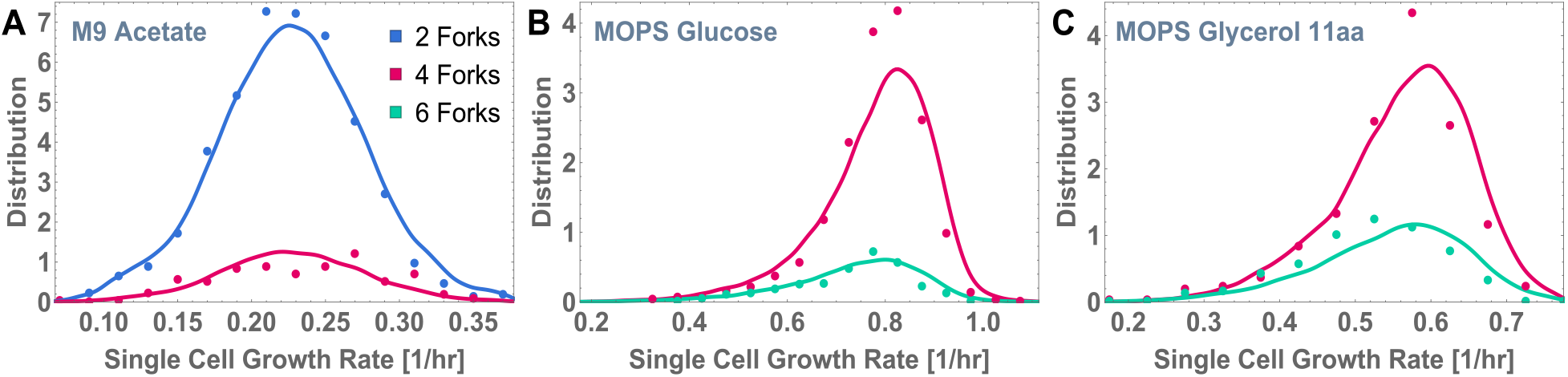
Comparing model predictions to experimental data. Model simulations (dashed lines) are compared with the experimentally obtained distributions (solid lines) of number of forks after initiation at different single-cell growth rates for the following growth conditions: (A) M9 acetate, (B) MOPS glucose and (C) MOPS glycerol 11aa. Colors represent different number of forks after initiation: 2 (blue), 4 (magenta), and 6 (teal). There is a close match between model predictions and experimental data, indicating that the presence of different numbers of forks and both single and multifork replication within the same growth condition is purely a consequence of the inherent stochasticity in the three time periods governing the replication cycle (the C period, the inter-initiation period, and the time to division), each of which depends solely on single-cell growth rate for a given growth condition.

Notably, although population MDTs in MOPS glucose and MOPS glycerol 11aa are on either side of the Cooper-Helmstetter 60-min MDT transition point between the slow and fast regimes (MDTs 52 min and 63 min, respectively), we did not observe an abrupt plateau in single-cell C period under either condition. In these conditions, single fork replication was entirely composed of the 4-fork replication pattern also seen in the slow, supposedly single fork regime (*τ_d_* > *τ_i_* > *τ_c_*), while multifork replication occurred when a new round of replication is initiated prior to completion of the ongoing round of replication, resulting in 6 forks (*τ_d_* > *τ_c_* > *τ_i_*). Thus, for these datasets, our framework requires only *τ_i_* and *τ_c_* to differentiate between single and multifork replication.

## Relationship between stochastic timers dictates replication fork number

In the CH model, and in the field in general, multifork replication is viewed as an adaption to the plateau in elongation rate observed in rapidly growing cells.^10^ While we do not observe a sharp or even a complete plateau in single-cell C period in any of the datasets, we still observe a clear divergence between single-cell growth rate and C period in fast-growing subpopulations suggesting that 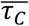 (the population averaged value), may eventually plateau under more nutrient-rich conditions.

Combining the behavior of our model’s timers during slow and intermediate growth with the available population-level data for fast growth conditions, we propose that the population average, 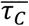, gradually flattens as *k* increases. This model is consistent with the observation that fork velocity varies as much as fourfold in single cells.^29^ From an unconstrained mathematical perspective of the population level, 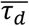 is expected to plateau as population MDT is reduced, since *τ_d_* is always greater than *τ_c_*, by definition. (**Fig. 3**) In contrast, due to the constraint that every division must be preceded by a corresponding initiation, 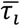 follows the same trend as MDT when the nutrient-imposed growth rate is varied.

Taken together, the measured and predicted timer behaviors suggest that multifork replication is a consequence of changes in the relationship between individual timers at fast single-cell growth rates. The timers differentially impact the relationships amongcell cycle events depending on growth regime (i.e., slow, intermediate, or fast growth). During slow growth, fork numbers are solely determined by the relative order of *τ_t_* and *τ_d_*, while during intermediate growth they are determined by *τ_t_* and *τ_c_*. During fast growth, we predict that all three timers play a role in determining fork numbers, especially during growth conditions that promote a mixture of allowable chromosome configurations and fork numbers. At the population level, our framework predicts the emergence of 8-fork (or more) cycles (*τ_i_* < *τ_c_* <2*τ_i_* < *τ_d_* <3*τ_i_*) during fast growth, a mixture of 4-fork (*τ_c_* <*τ_i_* < *τ_d_* <2 *τ_i_*) and 6-fork cycles (*τ_t_* <*τ_c_* < *τ_d_* <2 *τ_i_*) during intermediate growth (near 60 min MDT), and a combination of 2-fork and 4-fork cycles during slow growth. Supporting the validity of our framework, we obtained a close match with data from all three conditions without any fitting parameters (**Fig. 5B, C**), and an updated and more detailed rendering of the CH model (**Fig. 6** vs. **Fig. 1**).

**Fig. 6:**
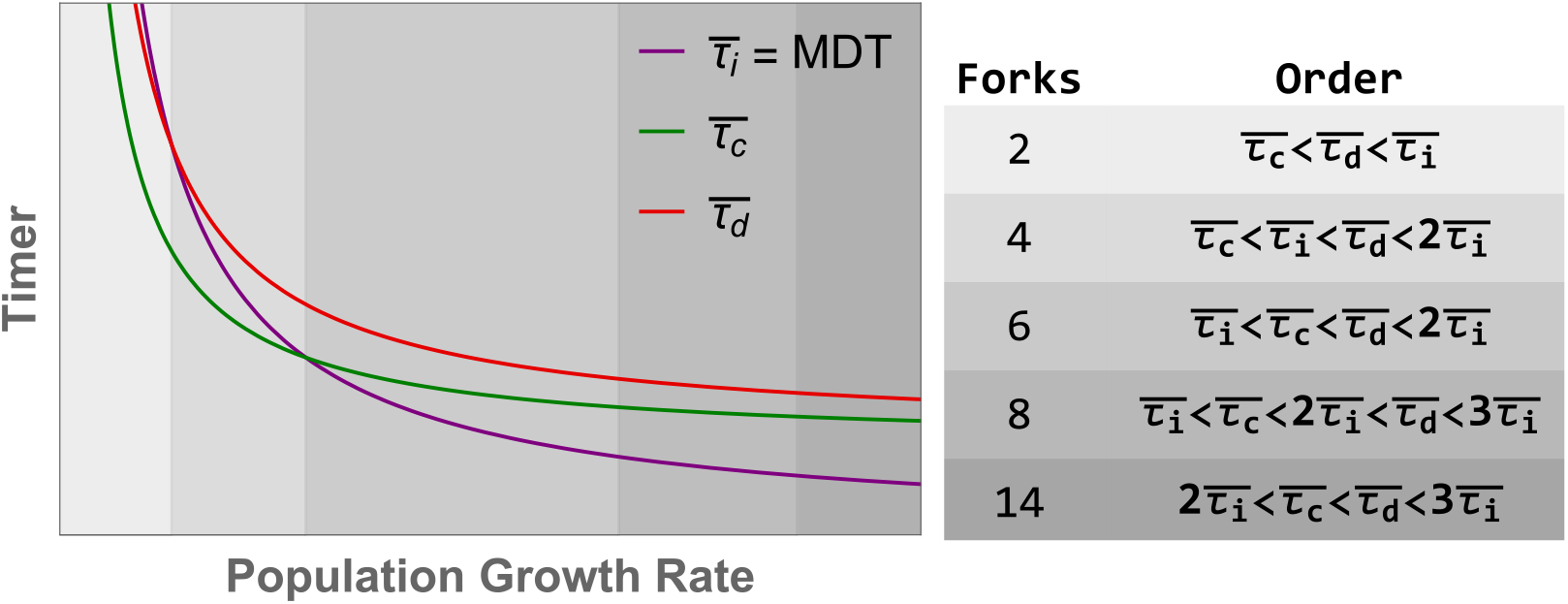
Determining dominant fork numbers from nutrient-imposed population growth rate. Our expectation for the trends in 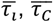, and 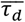 (representing the population mean values related to the stochastic timers *τ_i_*, *τ_c_*, and *τ_d_*) as a function of nutrient-imposed population growth rate. We expect mean *τ_c_* and *τ_d_* to flatten as growth rate goes to infinity, while mean *τ_i_* (equal to MDT) approaches zero. Fork numbers at different population growth rates are determined by the relative order of these timers, as shown.

## A roadmap for the application of this model to open biological questions

To test the ability of our agnostic framwork to address mechanism, we assessed the fork number-dependence of the divergent relationship between *τ_c_* and single-cell growth rate. The possible mechanisms responsible for divergence of *τ_c_* and growth rate are divisible into two broad mechanistic classes: replication fork-dependent and fork-independent. The distinguishing feature of these two classes is whether or not the number of forks present in a given cell impact replication speed. The traditional “maximum velocity” explanation falls under the fork-independent category. However, we do not favor this idea based on the fourfold difference in elongation rate observed in single cells under near steady-state conditions,^29^ and population data indicating that C period is reduced as much as 30% in short *E. coli* mutants.^36^

To distinguish between fork-dependent and independent models, we have plotted *τ_t_* and *τ_c_* relative to *k* in the two intermediate growth conditions and separate the population based on cycles containing either 4- or 6-fork cycles. (**Fig. 4**) Although a plethora of mechanisms could underlie fork number-related trends, we highlight two possibilities: titration and fork spacing models. A reduction in *τ_t_* and *τ_c_* (and consequently *τ_d_*) relative to *k* in a fast-growing population that is *independent* of total fork number could be consistent with titration of limiting replication substrates or enzymes (“titration model,” **Fig. 7A**). Conversely, reductions in *τ_t_* and *τ_c_* that are *correlated* with fork number could support a model in which physical constraints decrease the maximum replication rate due to the increased number of replication forks progressing on a single strand (“fork spacing model,” **Fig. 7B**). Applying this “test”, we observed diminishing returns to *τ_c_* as *k* increases, *independent* of the number of replication forks present in individual cells. (**Fig. 8**) Given the limited number of 4- and 6-fork cycles in the current dataset, our future work entails the full evaluation of this question with sufficiently large datasets representing a wider range of growth conditions. Then, we can fully dissect the mechanistic and molecular actors underlying this apparent fork-independence.

**Fig. 7:**
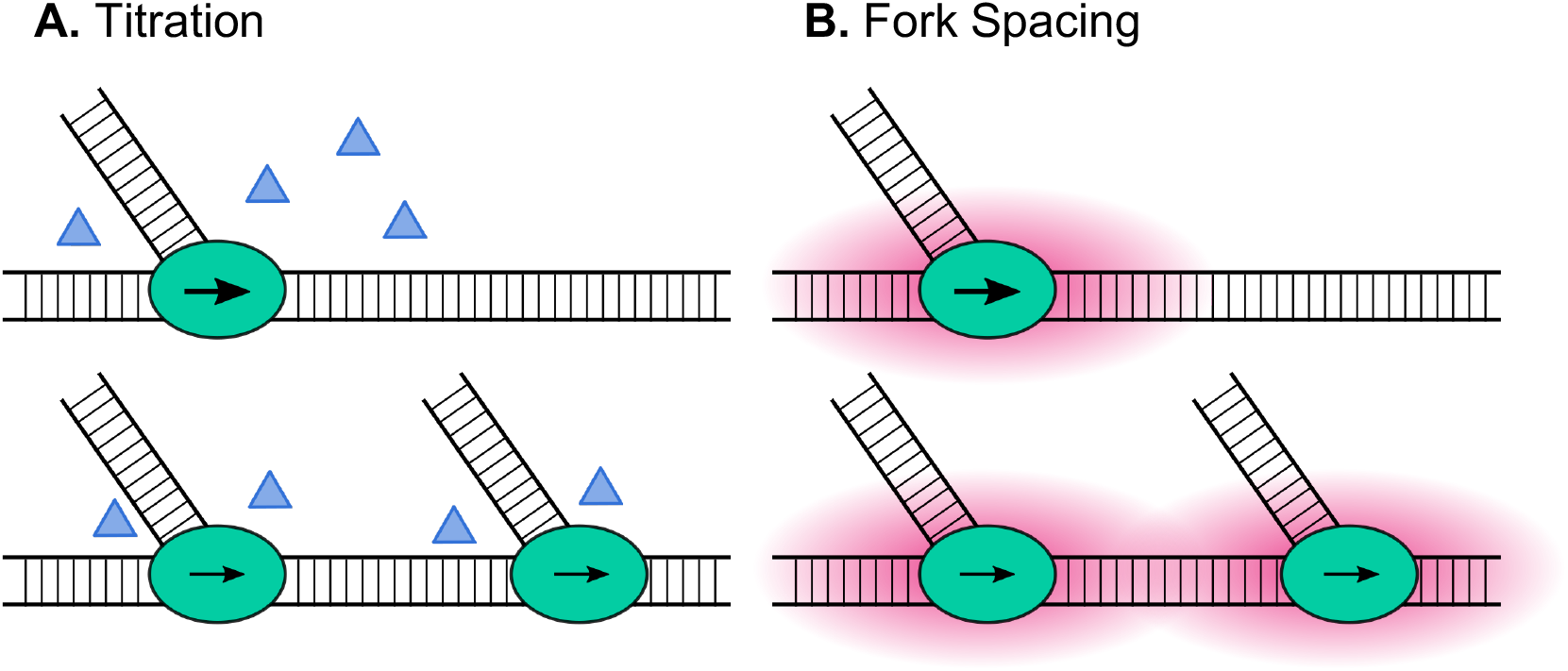
Two hypothesized models for C period decrease as growth rate increases. A) titration. B) fork spacing. Ovals represent active replisomes, triangles represent accessory replisome components and dNTP substrates, arrows represent relative replisome speed, and red clouds around the replisome represent steric repulsion and topological changes that alter replisome kinetics.

**Fig. 8:**
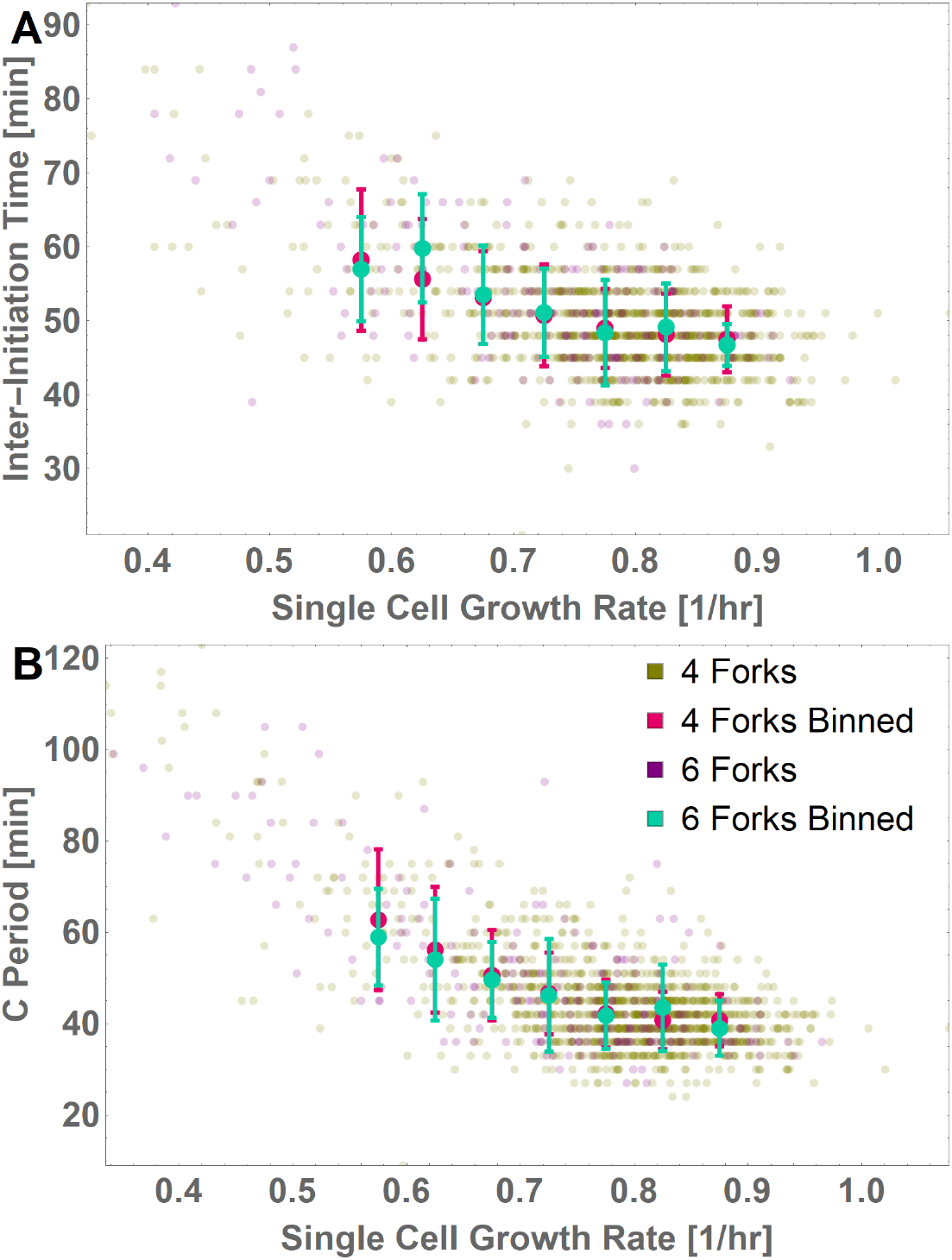
Both Inter-initiation time and C period are independent of fork number. (A) Inter-initiation time (*τ_i_*) and (B) C period (*τ_c_*) is plotted against single cell growth rate (*k*) for an intermediate growth condition (medium 4). Mean and S.D. of binned C periods and the spread of data points are plotted separately for 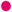 4-fork (single fork) and 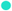 6- fork (multifork) data, based on the fork number observed just after initiation.

## Discussion

Scientific progress depends on our ability to incorporate new information, and is often driven by technological advancement. While the CH model has served a valuable function—contextualizing and inspiring work on the bacterial cell cycle for over 50 years—rapid advances in single-cell analysis reveal its limitations.

To fill this gap, we developed a new framework with which to understand cell cycle coordination. This framework offers numerous advantages for the evaluation of single-cell data. Importantly, it may be applied to any strain in any growth medium in any growth regime. Our framework is agnostic to the mechanism underlying the stochastic dynamics of initiation, replication and division, and simply captures these dynamics through the experimentally measured “calibration curves” for each of these timers as functions of single cell growth rate.s.

In sum, recognizing the value of single-cell data as a framework from which to understand the molecular mechanisms underlying cell cycle progression in bacterial cells is just the first step. Larger, more comprehensive single-cell datasets spanning a wide range of MDTs and media compositions is essential to determine the relationship between nutrient availability and cell cycle progression at high resolution not only in *E. coli* but in other bacteria, model and non-model alike. We look forward to the next chapter!

## Author Contributions

P.A.L. and S.I.-B. conceived of and designed research; S.S. and K.J. analyzed the extant data; S.S. spearheaded the writing process, contextualized the existing literature, and conceptually integrated biological and biophysical findings; K.J. formulated the model, analyzed the single-cell datasets, and performed analytic calculations and simulations under the guidance of S.I.-B.; all authors discussed the results and wrote the paper.

## Acknowledgments

We thank Suckjoon Jun and Fangwei Si for generously sharing the details of their methodology and analysis, along with the raw, single-cell datasets that are the heart of this study.^8^ We also thank them as well as Jade Wang for in-depth discussions and comments on the manuscript. We are grateful to the Levin and Iyer-Biswas groups for their insights and input as we pressure-tested multiple iterations of the model and its implications. P.A.L. and S.I.-B. thank the Aspen Center for Physics for graciously hosting us, and making our collaboration possible.

## Supplemental Information

**Fig. S1.**
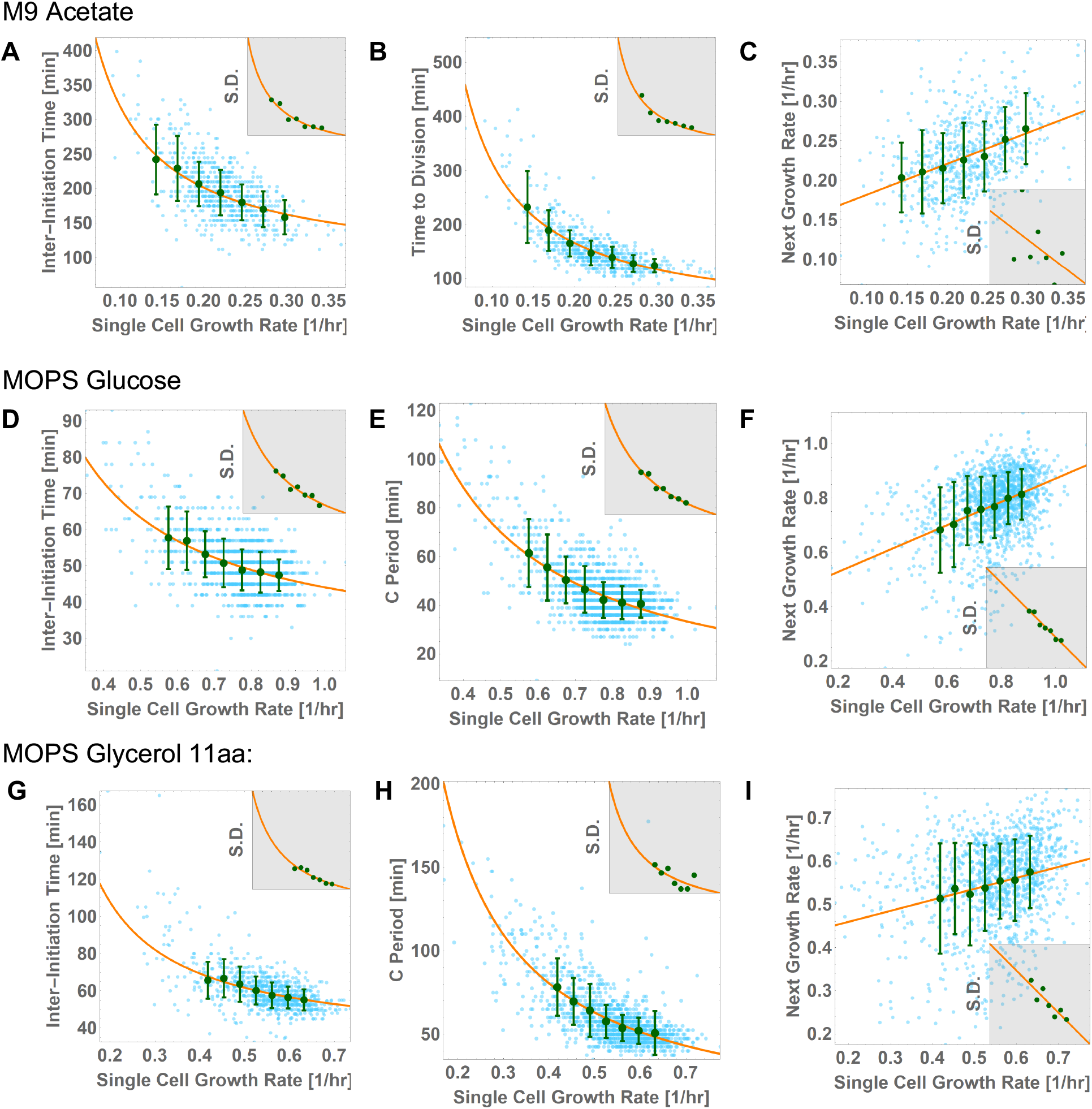
Calibration curves used as inputs in the model. For cells growing in (A-C) M9 Acetate, (D-F) MOPS Glucose, and (G-I) MOPS Glycerol 11aa, data points with binned mean and S.D. of (A, D, G) inter-initiation time, (B) Time to Division in slow growth condition and (E, H) C Period in intermediate growth conditions (as explained in the main text, intermediate growth conditions require C Period instead), and (C, F, I) single-cell growth rate of subsequent generation are plotted as a function of single-cell growth rate of current generation (inset: S.D. as a function of growth rate for the corresponding binned points). The orange line is the best-fit line (A and B: y=a+b/x and C: y=a+bx) fit using the binned mean. The observation that each binned mean and S.D. are quantitatively related enables us to use Eq. 1 to represent the different distributions at different single-cell growth rates within a growth condition by the same invariant rescaled distribution. This results in a significant simplification of model complexity. Orange best fit curves correspond to the mean and S.D. of these quantities, which we take as inputs to our model when using the rescaled distribution formulation; thus, their match with the green binned experimental data points is essential for accuracy of our results. Note that the S.D. in (C) inset for M9 Acetate is almost constant (with a variation of only 8%).

### Appendix on protocol for generating “calibration curves” from data

We tested our model on raw data published by the Jun lab (originally generated to study the validity of the independent double adder model of size control) ^8^ spanning three growth conditions: M9 acetate (MDT 195 min, equivalent to CH’s slow, single fork regime), and MOPS glucose and MOPS glycerol 11aa (MDTs 52 min and 63 min, respectively, both near the transition between CH’s single fork and multifork regimes).

To express the functional dependencies of the distributions of the three timers *τ_i_*, *τ_c_*, and *τ_d_* on single cell growth rate *k* in a form usable by our model, we identified emergent simplicities, or scaling collapses, and generated calibration curves for each media condition. Briefly, for each of the three growth conditions, we plotted all data points for the given condition relative to the singlecell exponential growth rate. We then subtracted the best-fit mean at the corresponding growth rate from each data point and divided the resulting value by the best-fit standard deviation (S.D.) at that growth rate. Finally, we measured the probability distribution of these scaled values, ***F***. Once again, the use of distinct calibration curves generated for each condition highlights the important relationship between medium composition and cell cycle dynamics, and reinforces previous work identifying the limitations of population “growth rate” as an independent variable with direct impacts on cell size and other phenotypes.^32–34^

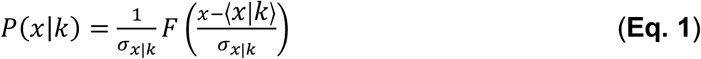

Within experimental error, the growth rate-independent distribution *F* can be appropriately scaled as shown in **Eq. 1** to give the conditional distribution of the stochastic timescale in question at any given growth rate, with the appropriate mean and variance, and approximate correct shape. Here, *x* is the stochastic timescale in question; *σ*_*x*|*k*_ and 〈*x*|*k*〉 are the fitted S.D. and mean, respectively, as functions of *k*. (See **Fig. S1 A, B, D, E, G, H**) We performed a similar analysis to characterize the dependence of *k* during a given inter-initiation cycle on its value during the previous cycle (**Figs. S1 C, F, I**). Together, these experiment-derived calibration curves serve as inputs in our model and are used in our simulations to generate the next *τ_i_*, *τ_c_*, *τ_d_*, and *k* for each consecutive replication cycle.

